# Design and implementation of a synthetic biomolecular concentration tracker

**DOI:** 10.1101/000448

**Authors:** Victoria Hsiao, Emmanuel LC de los Santos, Weston R. Whitaker, John E. Dueber, Richard M. Murray

## Abstract

As a field, synthetic biology strives to engineer increasingly complex artificial systems in living cells. Active feedback in closed loop systems offers a dynamic and adaptive way to ensure constant relative activity independent of intrinsic and extrinsic noise. In this work, we design, model, and implement a biomolecular concentration tracker, in which an output protein tracks the concentration of an input protein. Using synthetic scaffolds built from small, modular protein-protein interaction domains to colocalize a two-component system, the circuit design relies on a single negative feedback loop to modulate the production of the output protein. Using a combination of modeling and experimental work, we show that the circuit achieves real-time protein concentration tracking in *Escherichia coli* and that steady state outputs can be tuned.

## Introduction

Implementation of reliable feedback and control in engineered circuits is a continuing challenge in synthetic biology. Though positive and negative feedback systems are an essential feature of natural biological networks, synthetic circuits more commonly rely on library-based screening to find optimal expression levels. Not only are the resulting systems sensitive to relative concentrations between components, but each time the circuit is expanded, the network of regulatory sequences must be re-optimized to account for increased load on cell machinery (Klumpp *et al*, 2009). More importantly, this type of open loop approach only optimizes for a single set of environmental parameters, and inherently does not accommodate for stochastic cell-to-cell variation, changes due to cell growth cycles, or changes in cell loading from other circuit modules (Cardinale *et al*, 2013).

Closed loop systems provide robust regulation of individual components relatively independent of environmental disturbances. Negative feedback is a common feature of natural pathways, and has been shown to decrease transcriptional response time (Rosenfeld *et al*, 2002), to provide stability and reduce fluctuations (Becskei and Serrano, 2000), and to be necessary for oscillatory behavior (Ferrell, 2013). Given the stochastic and variable nature of protein production in the cell, regulation via relative concentrations ensures consistent protein ratios without having to rely on controlling absolute molecular counts.

Active feedback in biological systems has been previously considered at various levels. Recent studies have designed and studied an RNA-based rate regulating circuit with two opposing negative feedback loops (Franco *et al*, 2008), a system utilizing an RNA binding protein to repress translation of its own mRNA (Stapleton *et al*, 2012), and analysis of noise in transcriptional negative feedback (Dublanche *et al*, 2006). There have also been demonstrations of an *in silico* closed loop system, in which a computer measured fluorescence output and automatically modulated the activity of a photosensitive transcription factor (Milias-Argeitis *et al*, 2011). In that example, the negative feedback occurred in the software control system outside of the cell.

In this work, we present an *in vivo* protein concentration tracker circuit. To our best knowledge, this is the first application of synthetic scaffolds for real-time tracking of molecules entirely within the the cell environment at fast timescales. This circuit contains a single negative feedback loop implemented with synthetic scaffold proteins. We show that this feedback results in fast modulation of one protein concentration (the *anti-scaffold)* to track that of the reference protein (the *scaffold)* over a range of reference induction levels.

## Results

### Scaffold-based circuit design and implementation

Previously, Whitaker *et al* (2012) designed a scaffold-dependent two-component system in which the phosphotransfer was mediated by a synthetic scaffold protein consisting of small protein-protein binding domains. They demonstrated that weak natural cross-talk between a noncognate histidine kinase and response regulator pair could be artificially amplified via colocalization onto the scaffold. By fusing the kinase to the Crk SH3 domain and the response regulator to half of a leucine zipper, both would be recruited in the presence of a scaffold protein consisting of the SH3 ligand and the other half of the leucine zipper. Forcing the kinase and response regulator into close proximity greatly enhances the level of phosphotransfer. The kinase-regulator pair of Taz and CusR was chosen because of measured low levels of cross-talk upon long incubations of purified proteins (Skerker *et al*, 2005).

Building upon this scaffold-dependent two-component system, we designed a negative feedback circuit by introducing the anti-scaffold molecule. The *scaffold* molecule consists of a leucine zipper domain (LZX) linked to the SH3 ligand via flexible glycine-serine repeats (Figure 1). The two component system is comprised of the chimeric kinase Taz linked to four SH3 domains and the response regulator CusR linked to a single leucine zipper (LZx) domain (Figure 1A). The presence of the scaffold brings together the SH3 domain-ligand and LZX-LZx protein binding domains, recruiting Taz and CusR into close proximity and resulting in phosphorylation of CusR. The phosphorylated CusR becomes an active transcription factor, binding to its natural promoter (P_CusR_) and activating expression of the *anti-scaffold* protein (Figure 1B). The anti-scaffold consists of the complementary LZx and SH3 ligand domains, which allow it to competitively bind to and consequently sequester the scaffold protein (*K_d_* = 6 nM for the leucine zipper and *K_d_* = 100 nM for the SH3 domain (Acharya *et al*, 2002; Whitaker and Dueber, 2011)). This prevents further phosphorylation of the response regulator, and halts further production of the anti-scaffold. In the absence of any scaffold protein, no activated response regulator activity is observed (Supplementary Fig. S2).

**Figure 1:**
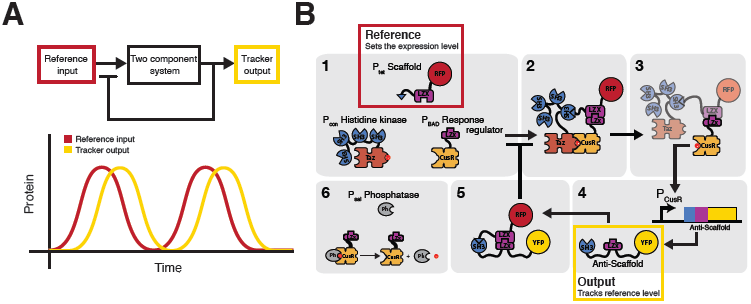
Overview of circuit design. A) The circuit takes an input that sets the reference value. The input proportionally modulates activity of a two component signaling system that then produces an output. The output triggers a negative feedback response. The desired circuit function would be real-time tracking of the input by the output. B) The specific implementation of the circuit is shown. The circuit regulates the production of the amount of target protein (Anti-scaffold-YFP) with respect to the amount of reference protein (Scaffold-RFP). Expression of the target is dependent on the amount of free scaffold. The target contains domains which sequester free scaffold creating a negative feedback loop. Scaffold, response regulator, and phosphatase concentrations are induced via P_tet_, P_BAD_, and P_sal_, respectively.

We implemented the circuit in a ΔCusS ΔCusR *E. coli* knockout strain (Whitaker *et al*, 2012). In the absence of CusS, the native bifunctional histidine kinase/phosphatase partner for CusR, activated CusR proteins remain phosphorylated indefinitely. Accordingly, we re-introduced a CusS(G448A) mutant behind an inducible promoter to control response regulator deactivation. The G448A mutation disrupts the ATP binding site, eliminating kinase autophosphorylation without affecting phosphatase activity (Whitaker, 2012; Zhu and Inouye, 2002). This created a tunable phosphate sink in our circuit and ensures tight coupling between present scaffold and activated response regulator concentrations. The negative feedback circuit with the anti-scaffold is referred to as the *closed loop circuit*. As a control, we also built an open loop circuit, which instead of P_CusR_-driven expression of the anti-scaffold, only YFP is expressed.

We constructed the circuit as a three plasmid system, in which the kinase is constitutively expressed and the scaffold, response regulator, and phosphatase were cloned behind the inducible promoters P_tet_, P_BAD_, and P_sal_, respectively. We created scaffold-RFP and anti-scaffold-YFP fusion proteins to track temporal changes in concentrations *in vivo*. The fluorescent reporters mCherry RFP and Venus YFP were chosen on account of their similar maturation times (approximately 15 min) (Nagai *et al*, 2002; Shaner *et al*, 2004). To visualize dynamic behavior, the scaffold-RFP and anti-scaffold-YFP fusions were tagged with a C-terminal ssrA degradation marker of medium strength (RPAANDENYAAAV) (Andersen and Molin, 1998).

### Modeling dynamics and steady state circuit behavior

Building upon a previously published model of this basic scaffold-modulated circuit, we created additional phosphatase species and reactions (de los Santos *et al*, 2013). The model is differential equation based and all chemical reactions between species are explicitly stated, omitting transcriptional activity and accounting only for protein level behavior. With the exception of the anti-scaffold production term, all other terms are derived from mass action kinetics. The 25 species arise from combinations of scaffold (Sc), response regulator (RR), histidine kinase (HK), anti-scaffold (AS), and phosphatase (Ph) binding complexes. In total, the model consists of 80 reactions, 25 differential equations, and 26 parameters (See Supplementary for complete list of chemical reactions). Many parameters (Table 1) were selected from experimental values found in the literature (Pazy *et al*, 2009; Groban *et al*, 2009; Solomaha *et al*, 2005), and others were explored within a physiologically reasonable range.

**Table 1:**
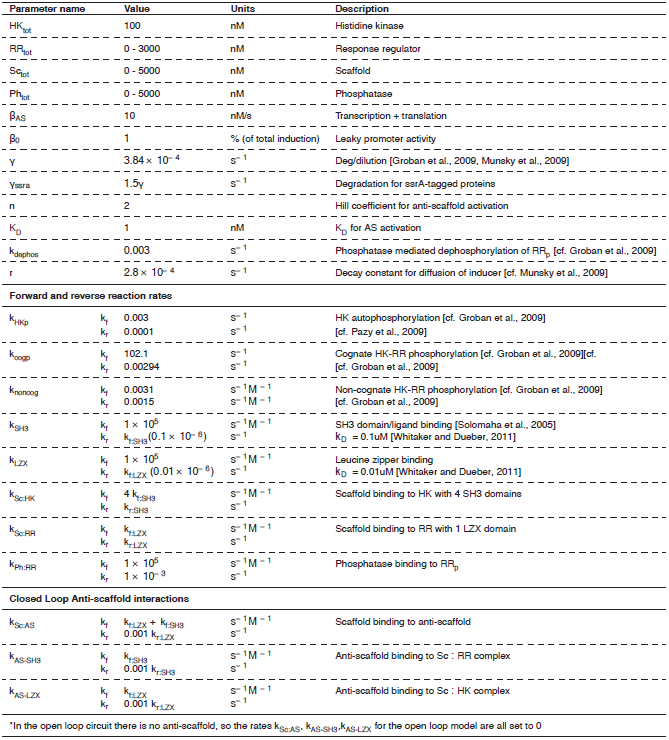
Table of model parameters estimated from literature.

Model reactions can be classified into five categories: production and degradation, phosphorylation, scaffold complex formation, activation, and irreversible sequestration. Phosphorylated species are denoted with a subscript p (e.g. RR_p_), and complexes are denoted with a colon separating the participating species (e.g. Sc:AS). Though the possibility of modeling the scaffold as an enzyme-like species was considered, we could not assume that either the kinase or response regulator would always be in excess, a requirement of the substrate in a Michaelis-Menten reaction. Therefore, Michaelis-Menten kinetics were deliberately avoided.

The production rate, *β*, of the scaffold, histidine kinase, response regulator, and phosphatase are determined by user input of the total steady state value (in nM), multiplied by the degradation/dilution rate *γ*. This ensures constant concentration of these species in solution. The degradation rate *γ* is applied universally for all species and is estimated based on a cell division time of 30 minutes (Groban *et al*, 2009). Experimentally, the scaffold is on a high copy plasmid, the response regulator is on a medium copy plasmid, and the kinase and phosphatase are on the same low copy plasmid.

The phosphorylation reactions describe the autophosphorylation of HK and dephosphorylation of RR_p_. Key reactions that describe this process are:

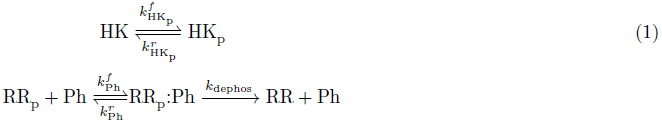

The phosphatase forms a complex with the RR_p_ prior to dephosphorylation. We model both phosphorylation and dephosphoryaltion with a two-step reaction model, an approach consistent with previous models (Huang and Ferrell, 1996). Rate constants for kinase phosphorylation and dephosphorylation of the response regulator were chosen based on cognate and noncognate phosphorylation rates measured for natural two-component systems, and occur on the order of seconds (Groban *et al*, 2009). The following equations show phosphorylation in the absence and presence of scaffold:

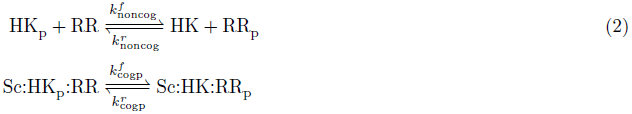

Reaction rates for scaffold complex formation were based on the kinetics of the protein-protein interaction domains SH3 domain/ligand and LZX/LZx. SH3 domain/ligand binding has an estimated association affinity *K_d_* of 0.1 uM while leucine zippers have a *K_d_* of approximately 0.01 uM (Posern *et al*, 1998; Acharya *et al*, 2002; Grunberg *et al*, 2010; Whitaker and Dueber, 2011). Here we have examples of histidine kinase and response regulator binding to scaffold via SH3 and LZX binding, respectively:

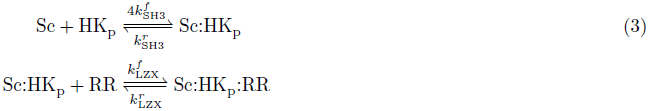

A phosphorylated response regulator becomes an active transcription factor. We considered all possible complexes with RR_p_ as possible activators. This sum of all possible species is referred to as RR_active_. The response regulator, CusR, dimerizes upon phosphorylation, so we use a Hill function with coefficient *n* = 2 to represent activation of AS production:

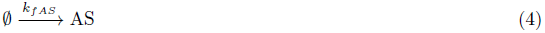

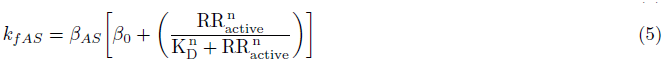

where RR_active_ = RR_p_ + Sc:RR_p_ + Sc:HK:RR_p_ + Sc:HK_p_:RR_p_ + Sc:RR_p_:AS.

The negative feedback component comes about through the irreversible sequestration of the scaffold once it has bound to the anti-scaffold. We made the assumption that the individual SH3 and LZX domains on the anti-scaffold bind independently, at the same rates as HK and RR binding. However, once either the SH3 or LZX component of the AS has bound to the Sc, resulting in a local concentration of the free domain that is substantially higher than the *K_D_*, the other domain quickly displaces any competing species and sequesters the entire Sc. The effective irreversibility comes about through steric hinderance of competing HK and RR species, both of which only have one compatible binding domain to the Sc:

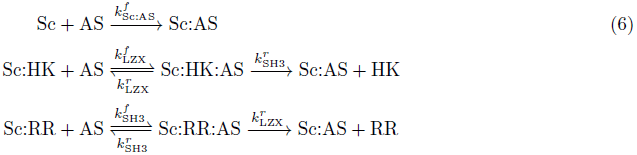

After creating the model, we tested its validity by comparing the open and closed loop circuits. In the open loop circuit, the negative feedback binding reactions are set to zero (Table 1). Experimentally, this was done by replacing the anti-scaffold with a fluorescent reporter alone. Figure 2A shows simulated steady state values for anti-scaffold (or fluorescent reporter) output over a range of scaffold concentrations (0 - 1000 nM), with either 0 nM or 100 nM of response regulator. In the cases with no response regulator, the circuit does not function and production of output is solely doe to simulated leaky anti-scaffold production (*β*_0_). When response regulator molecules are present, the open loop circuit output decreases significantly with increasing scaffold. Though it is not intuitive, this can be explained as the scaffold single occupancy effect (Whitaker and Dueber, 2011; Good *et al*, 2011), where an overabundance of free scaffold leads to binding of only kinase or response regulator but not both. As we examine the prevalence of these intermediate species (Sc:HK, Sc:RR) in simulation, we can see that the total concentration of singly-bound scaffold increases, decrease in output is indeed observed(Supplementary Fig. S1A). The same effect also occurs in the closed loop circuit, but much higher concentrations of scaffold are needed, since the anti-scaffold sequestration lowers the effective number of free scaffold molecules in solution (Supplementary Fig. S1B).

**Figure 2:**
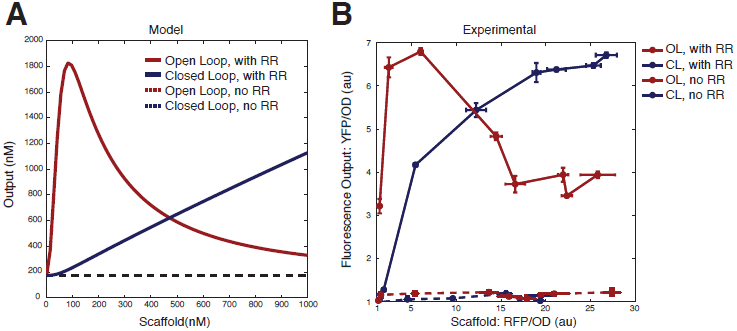
Open loop versus closed loop. A) Model predictions of scaffold circuit with and without negative feedback. Solid lines show anti-scaffold output over a range of scaffold concentrations (0 - 1000 nM) for open and closed loop circuits with constant response regulator (100 nM). Dotted lines show lack of output in the absence of response regulator. Open loop circuit shows scaffold single occupancy effect. B) Steady state experimental data of open and closed loop circuits with and without response regulator matches model predictions. Both sets of experimental data were normalized by the autofluorescence of the *E. coli* strain.

Experimental data for the circuit closely recapitulated the model predictions (Figure 2B). First, without induction of RR, there is no output YFP. Secondly, we see the single scaffold occupancy effect in the open loop circuit, but not the closed loop circuit. In the case of no scaffold induction, the open loop circuit has about three times more background than the closed loop circuit (Figure 2A), behavior that is predicted by the model (Figure 2B). We believe this is due to leakiness in scaffold production in the absence of aTc. In the closed loop circuit, this leaky production is subdued by the negative feedback, while in the open loop, we see significant production of YFP. The open and closed loop circuits have the same maximum output, but only the closed loop circuit shows a linear increase with increasing scaffold. These experiments validated our model, and demonstrated that this synthetic scaffold technology could be used for negative feedback.

### Model-informed exploration of parameter space

Once we validated the model, circuit limitations were explored *in silico*. Specifically, we investigated the effects of tuning response regulator and phosphatase concentrations, parameters which were accessible via inducible promoters in our experimental system. In Figure 3A, a scan of input-output response curves is shown over a range of response regulator and phosphatase concentrations. For each curve in the grid, the scaffold concentration in which the single occupancy drop-off occurs was found, and the slope of the curve up to that concentration was found with a linear fit. The maximum scaffold occupancy limit is the concentration of scaffold molecules at which each scaffold molecule only has either a response regulator or histidine kinase. The slope of the curve up to that point represents the anti-scaffold to scaffold ratio which can be achieved by the circuit. In the case where the single occupancy limit does not appear, the last concentration is used. Data shown in Figure 3B indicates that increasing response regulator values result in a greater AS/Sc ratio (up to 50% fold increase), while increasing phosphatase serves to bring down that ratio. A majority of the space tested shows that AS/Sc ratio is approximately 1:1 (See Figure S3 for explicit values). The effect of increasing phosphatase is apparent when the maximum scaffold occupancy limit is examined (Figure 3C). As phosphatase concentration increases, active response regulators are quickly dephosphorylated, decreasing the efficacy of the scaffolds, lowering the maximum occupancy concentration, and making the drop-off more steep.

**Figure 3:**
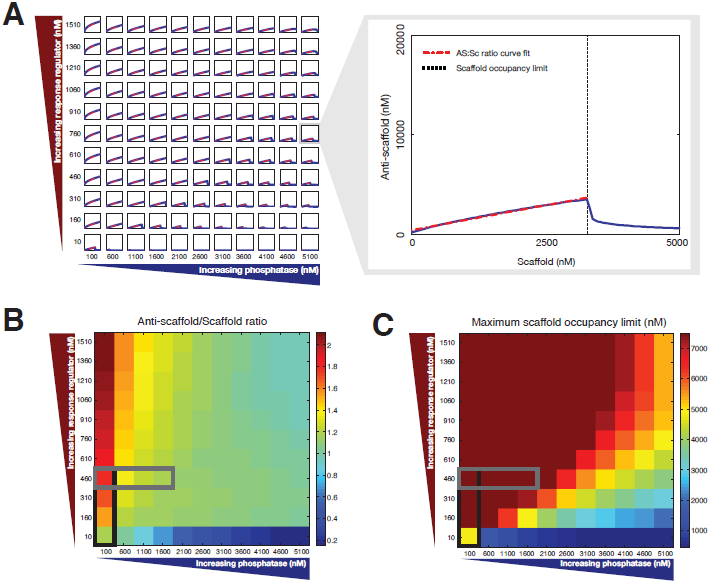
Model-based exploration of parameter space. A) Simulations of scaffold to anti-scaffold inputs and outputs over a range of phosphatase (100 - 5100 nM, 500 nM increments) and response regulator (10 - 1510 nM, 150 nM increments) concentrations. Enlargement shows the scaffold single occupancy limit concentration and curve fitting for each curve. Red dotted lines show curve fits - the slope represents the anti-scaffold to scaffold ratio. B) Heat map showing anti-scaffold to scaffold ratio for each curve shown in part A. Increasing response regulator results in greater AS/Sc ratios. Gray box represents estimated experimental phosphatase induction range. Black box estimates experimental response regulator induction range. C) Heat map of maximum scaffold occupancy limit. Higher concentrations of phosphatase result in decreased maximum scaffold occupancy limit.

By modulating response regulator and phosphatase concentrations, a range of maximal expression levels for scaffold and anti-scaffold can be achieved. Figures 4A and C shows steady state circuit response to varying levels of response regulator induction in both the model and experimental circuit. Increasing RR concentrations increases the gain of the system by increasing the number of available active transcription factors for the AS promoter. In simulation data (Figure 4A), we see that the scaffold occupancy effect is mitigated by higher levels of response regulator. This is consistent with our previous explanation, since more regulator means almost all free scaffold molecules will exist as Sc:RR. Experimental data for tuning response regulator concentration via ten-fold increases of arabinose (Figure 4C) do not extend the scaffold levels far enough to show the occupancy effect, but the increasing output gain is evident.

**Figure 4:**
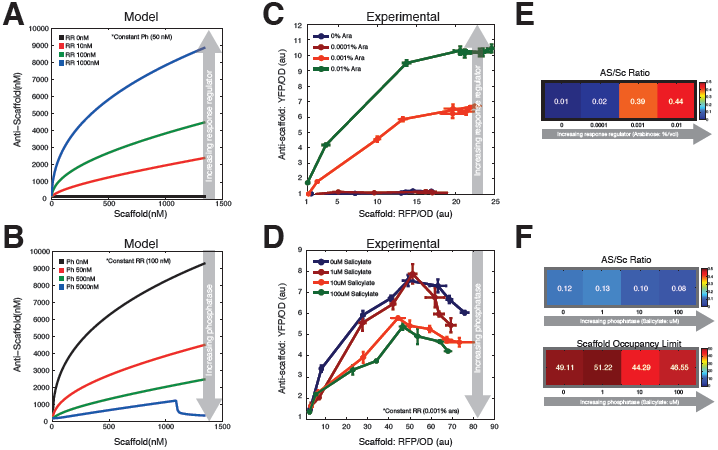
Steady state experimental tuning of response regulator and phosphatase. A) Simulation data of input-output curves with increasing response regulator concentrations (0 - 1000 nM). Increasing response regulator increases the scaffold occupancy limit as well as overall AS/Sc ratio. B) Simulation data of input-output curves with increasing phosphatase concentrations (0 - 5000 nM) with constant response regulator concentration of 100 nM. Increasing phosphatase decreases the scaffold occupancy limit and overall AS/Sc ratio. C) Experimental data of steady state scaffold to anti-scaffold curves with ten-fold increases in response regulator induction (0 - 0.01% arabinose). D) Experimental data of steady state circuit behavior with ten-fold increases in salicylate. Response regulator concentration is constant (0 - 100 uM salicylate). E) Heat maps showing quantified AS/Sc ratio and scaffold occupancy limit values with increasing response regulator. Scaffold occupancy limit was not observed in response regulator experiments. F) Heat maps showing quantified AS/Sc ratio and scaffold occupancy limit values with increasing phosphatase. All experimental data was normalized by baseline auto-fluorescence values.

The presence of phosphatase in the circuit modulates the amount of time that phosphorylated response regulator is active. Hence, tuning phosphatase concentrations changes RR ↔ RR_p_ cycling time. Figures 4B and D shows steady state responses across a range of phosphatase concentrations. Simulation results show that increasing phosphatase decreases overall circuit output (Figure 4B) by decreasing the average time RR_p_ is active. Experimental results (Figure 4D) support model predictions and show this suppression of output with increased induction via salicylate.

In Figures 4E and F, these experimental steady state data are analyzed using the same techniques shown in Figure 3. Figure 4E shows anti-scaffold to scaffold ratio and scaffold occupancy limit as calculated based on fluorescence data with ten-fold increases in response regulator induction with no phosphatase present. Similar to the analysis used in the model, if the single occupancy drop is not observed, the highest scaffold concentration is taken. Figure 4F shows the same metrics with ten-fold increases in phosphatase induction with constant response regulator (0.001% arabinose induction). Experimental data is presented as a function of fold change from background fluorescence, and so cannot be compared directly with model data (presented in nM). However, the overall trends are in agreement. As response regulator increases, we see a significant increase in anti-scaffold to scaffold ratio, and little change in the occupancy limit. With increasing phosphatase, we see a slight decrease in AS/Sc ratio and scaffold occupancy limit. We believe these data show us that our experimental range occupies only a small fraction of that shown by our model (Figure 3B,C), and that these limitations are due to the limited dynamic range of the P_BAD_ and P_sal_ inducible promoters (P_BAD_-RR, P_sal_-Phos).

### Characterization of step response

Having verified the functionality and potential range of the negative feedback loop, we tested circuit response to step inputs. Using a microfluidic plate under a microscope, step induction of the scaffold protein was achieved by flowing in 0, 37.5, or 75 nM of aTc after 30 minutes of growth in normal media (Figure 5). Cell production of response regulator and phosphatase was pre-induced by incubating cells with arabinose and salicylate. In all conditions, expression of scaffold-RFP began about 30 minutes after induction, and occurred almost simultaneously with that of anti-scaffold-YFP. In order to better visualize the fold change, fluorescence output is normalized by the maximum value of the lowest step input.

**Figure 5:**
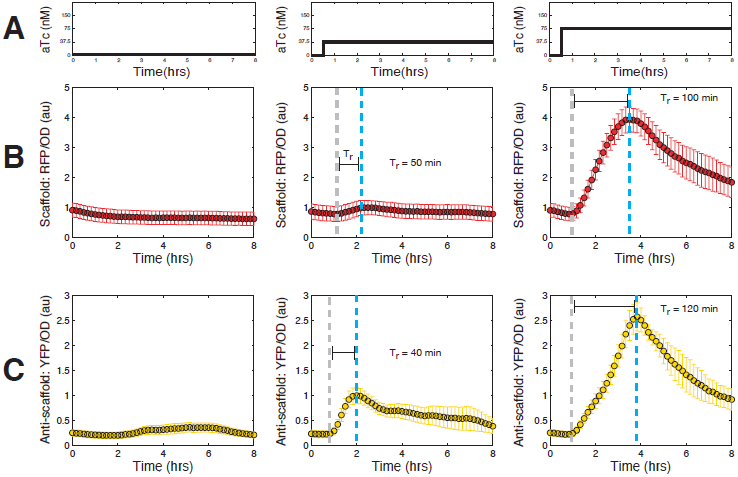
Step induction of closed loop circuit. A) aTc induction of Sc-RFP began 30 minutes after start of experiment and continued for the rest of the experiment. B) Scaffold-RFP/OD measurements for no induction (left), 37.5 nM induction (middle), and 75 nM induction (right). Response time (*T_r_*) is quantified by finding the time needed for fluorescence to increase from 10% (gray dotted line) to 90% of the maximum value (blue dotted line). A two-fold increase in aTc results in a four-fold increase in scaffold expression and a two-fold increase in response time. C) AS-YFP/OD measurements show 2.5 fold increase between the two inputs and a three-fold increase in response time. Fluorescent measurements are normalized such that the maximum of the middle column (37.5nM aTc) is 1 a.u. to better visualize fold change.

We quantified the response time of the circuit by calculating the response time (*T_r_*) of scaffold (RFP) and anti-scaffold (YFP). In control theory, response time is the amount of time needed for an output signal to increase from 10% to 90% of its final steady state. Here, we use the maximum output rather than a final steady state. As cells reach stationary phase, circuit expression gradually turns off, and no steady state in fluorescence output is maintained. Figure 5A shows that scaffold induction, regulated by a P_tet_ promoter, has a 4-fold expression increase between 37.5 nM and 75 nM induction, but only a 2-fold increase in response time (50 min to 100 min). In Figure 5B, we see that anti-scaffold output, regulated by the scaffold concentration, shows a 2.5 fold increase in maximum expression and a 3-fold increase in response time (40 min to 120 min).

This step input characterization revealed that scaffold and anti-scaffold fluorescence could be observed almost simultaneously about one cell cycle (30 min) after aTc induction of scaffold transcription. Following induction of the circuit, the response time to maximum expression increases in a linear-like fashion with increasing scaffold induction.

### Circuit closely follows three step induction

Following step input characterization, we investigated circuit response to multiple step-up inputs. Figure 6 shows the results of a three step scaffold induction experiment with one hour steps corresponding to 50nM increases of aTc inducer. The single negative feedback loop in the circuit represses overproduction of anti-scaffold but there is no mechanism for feedback in the case of an excess of scaffold or anti-scaffold. As such, the model predicts that increases in inducer will lead to immediate increases of scaffold followed closely by the anti-scaffold but once induction is turned off, degradation of proteins depends on the endogenous ClpXP degradation machinery (Figure 6B). Additionally, the upward slope of each curve should overlap until induction ceases.

**Figure 6:**
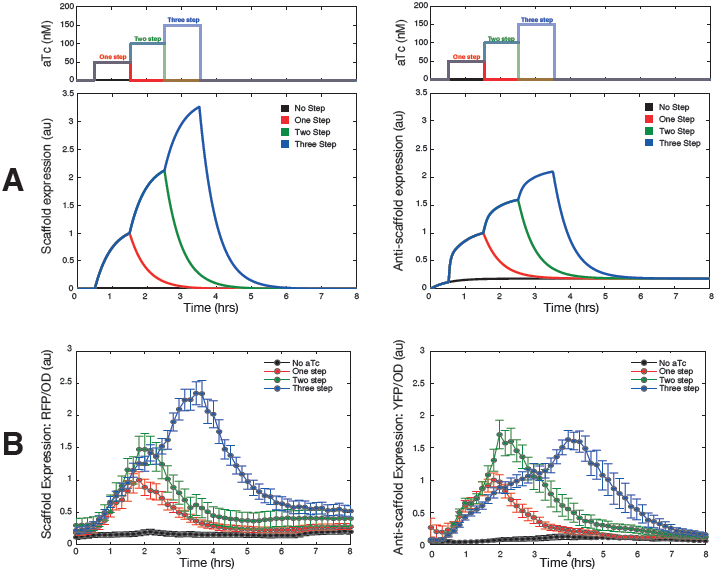
Multi-step induction of tracker circuit. A) Simulation results for a three step induction show overlapping response times with each curve decreasing based on degradation rate after induction ceases. Upper panel shows aTc induction pattern with one hour steps increasing in 50 nM increments starting 30 minutes after start of experiment. B) Experimental time traces for Sc-RFP show overlapping fluorescence output, with each curve decreasing at a time proportional to the number of steps. Corresponding anti-scaffold-YFP data show similar overlaps and proportional decreases. Fluorescent measurements are normalized such that the maximum value of the one step curve is 1 a.u. to better visualize fold change.

Step-up induction was performed on cells pre-incubated in arabinose and salicylate, activating expression of response regulator and phosphatase, respectively. As shown in Figure 6C, experimental results for a three step induction are consistent with model predictions, and show overlapping curves during the ascent, with each individual curve dropping off slowly as induction ceases. The chemical induction of the scaffold produces a much smoother output curve compared to the response regulator-modulated anti-scaffold.

### Inducer diffusion rates contribute to cumulative effect of sequential pulses

We observed in our model that variations in inducer diffusion rate would greatly affect the outcome of sequential pulses (Figure 7A). The removal of aTc from the cytoplasm and surround media is not instantaneous, and induction does not go to zero. Given two sequential 30 minutes pulses spaced one hour apart, the diffusion constant determined whether two independent, identical outputs occurred, or if an additive effect would take place. Essentially, if the first pulse of inducer is not given sufficient time to diffusion out of environment, aTc molecules from the first pulse are still present when the second pulse occurs. We modeled inducer diffusion following a pulse with an exponential decay term, *β_Sc_* = *β_ind_* exp(*−rt*) (Munsky *et al*, 2009). Figure 7A shows two pulse simulation results when the default decay constant (*r* = 2.8 × 10^−4^/s, middle column) is increased or decreased by 10-fold.

**Figure 7:**
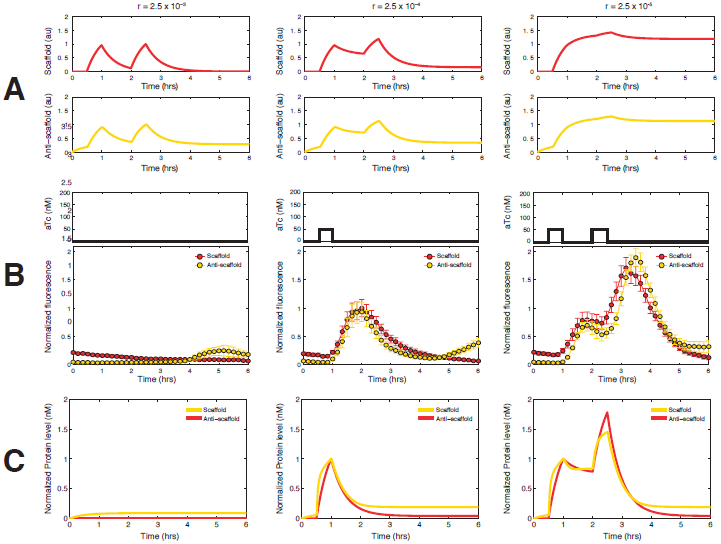
Two pulse induction of circuit. A) Model results for a range of inducer decay constants from 2.8 × 10^−3^ to 10^−5^/s. Fast diffusion (left) shows two independent pulses, intermediate diffusion (middle) results show some overlapping protein from first and second pulses, and slow diffusion (right) shows large amounts of overlapping protein from the first to the second pulse. B) Experimental data for zero, one, and two pulses of 50 nM aTc. Data are normalized by maximum of single pulse induction (middle column). C) Simulations with improved inducer diffusion rates.

When we tested two pulse induction *in vivo* (Figure 7B), we ran simultaneous experiments with zero, one, and two 30 minute pulses of 50 nM aTc. The single pulse fluorescence maximum (Fig. 7B, middle column) was normalized to 1 a.u. It is clear from the two pulse fluorescence output data that the diffusion rate of aTc after a pulse *in vivo* was actually much slower than expected *in silico*. In fact, so much of the scaffold from the first pulse remained that there was almost a two fold increase in maximal expression during the second pulse. This was an effect which had not been apparent previously during the multi-step inductions, where we showed sequential increases in inducer concentration. This set of data shows that modulation of pulse frequency, but not concentration, can result in the same additive effect as increasing inducer concentration.

We then sought to improve our model by tuning the inducer decay constant (Figure 7C), generating outputs which demonstrated the nearly two-fold increase observed *in vivo*. Although the slower rate better captured gene expression during log phase, we consistently observed a rapid decrease in fluorescence as cells approached stationary phase. We believe this is due to upregulation of ClpX and other ssrA machinery in stationary phase (Farrell *et al*, 2005). This resulted in improved model performance when simulating dynamic circuit behavior.

## Discussion

We have designed a novel negative feedback tracker circuit using modular synthetic scaffold proteins and a two-component system with scaffold-dependent phosphorylation. The use of scaffold proteins for negative feedback could potentially be a robust way of linking modules and ensuring constant performance despite intrinsic and extrinsic noise. Scaffold proteins have been shown to be powerful hubs for organization of regulatory feedback in natural networks, usually by colocalization of phosphorylation machinery (Good *et al*, 2011). Previous studies have rewired the naturally occurring Ste5 scaffold in the yeast MAPK cascade to redirect signals, to modify delays in signaling time, and to introduce ultra sensitivity (Park *et al*, 2003; Bashor *et al*, 2008). Here we have taken synthetic scaffold proteins, which were designed and optimized to increase flux through metabolic pathways (Dueber *et al*, 2009; Moon *et al*, 2010; Whitaker and Dueber, 2011), and built an entirely synthetic feedback circuit. The system is designed with multiple inducible promoters and most components can be tuned. Most importantly, the qualitative circuit behavior of anti-scaffold proportionally tracking the scaffold is maintained over a range of component concentrations.

After we designed the circuit framework, we constructed and then experimentally validated an ODE-based mathematical model. Through selection of parameters and reaction rates based on the literature, we obtained a model able to reasonably predict circuit behavior. Comparisons between simulation and experimental data confirmed the presence of scaffold-mediated negative feedback, and we used the model to scan the parameter space in a way that would have been time and resource intensive to explore *in vivo*. We found that steady state circuit gain can be tuned by changing response regulator concentrations and cycling time is controlled by varying phosphatase levels, observations which were supported by experimental data. Following initial step induction system characterization of step input response time, expression of both the reference (Sc-RFP) and output (AS-YFP) protein was shown to be fast and responsive to multi-step inputs. Finally, we found that pulse-modulated induction could result in additive circuit response, leading to improvement of the model through more accurate inducer diffusion parameter values.

This work demonstrates the design and implementation of a scaffold-based biomolecular tracking circuit that has potential applications in active regulation of component expression in synthetic circuits. The relatively small size (approx. 60 AA) of the scaffold and anti-scaffold proteins facilitates attachment to larger proteins, represented in this work by mCherry-RFP and Venus-YFP. Rather than open loop tuning of regulatory sequences and large-scale screening, scaffold-based negative feedback could be utilized. By attaching the scaffold to a native protein, it may also be possible to tie synthetic circuit inputs to naturally occurring cycles *in vivo*. It is well known that many natural cell processes such as developmental segmentation, circadian clocks, and stem cell multipotency involve oscillatory gene expression (Bessho, 2003; Imayoshi *et al*, 2013).

Furthermore, response to signal transduction may be modulated not by amplitude, but by frequency (Cai *et al*, 2008). We have shown that the scaffold-modulated protein tracker follows changes in both amplitude and frequency, and exhibits good agreement with a mass-action model. Future iterations of this design may improve tracking fidelity by including reverse feedback loop to compensate for overexpression.

## Materials and methods

### Cell strain, media

The circuit was implemented in the *E.coli* cell strain WW62, a variant of BW27783 (CGSC 12119) with knockouts of EnvZ, OmpR, CusS, CusR, CpxA, and CpxR. All cell culture was done in optically clear MOPS EZ Rich defined medium (Teknova, M2105), with 0.4% glycerol instead of 0.2% glucose. The use of glycerol as a carbon source was done to prevent interference with the arabinose induction of the P_BAD_ promoter.

Tested arabinose induction levels were 0, 0.0001%, 0.001%, 0.01%, and 0.1% (20% stock solution). Anhydrotetracycline (aTc) was diluted in media at concentrations of 0, 5, 15, 30, 60, 90, 120, 150 nM. Sodium salicylate was resuspended at a stock concentration of 100 mM and diluted 1:1000 in media for experiments.

### Plasmids

The plasmid encoding the SH3-ligand-LZX-mCherry scaffold (pVH001) has a high copy backbone(ColE1) with ampicillin resistance. The CusR-LZx response regulator and SH3-domain-LZx-VenusYFP anti-scaffold plasmids (pVH003 for closed loop, pVH009 for open loop) are on a medium copy backbone (pBBR1) with kanamycin resistance. The 4SH3-domain-Taz histidine kinase and CusS-G448A phosphatase are on a low copy plasmid (p15A) with chloramphenicol resistance. Detailed plasmid maps are shown in Figure S4, and a complete list of plasmids and strains can also be found in the Supplementary Information.

### Microscopy

Step induction data were taken using the CellAsic ONIX microfluidic perfusion system for bacteria (B04A). The microscope is an Olympus IX81-ZDC enclosed in a custom heater box. Images were taken using a 100x oil immersion phase objective. Fluorescence filters are 580/630 for mCherry (Chroma 41027) and 510/560 for YFP (Chroma 31040 JP2). Microscope media was augmented with oxidative scavengers Trolox (200 nM) and sodium ascorbate (2 mM).

Cells are pre-induced with arabinose(0.01%) and salicylate(100 uM) to ensure the RR and Ph are pre-expressed prior to addition of aTc. Overnight cultures are diluted 1:500 in media containing arabinose and/or salicylate four hours prior to loading in the CellAsic plate. Cells are diluted 1:10 again before loading. During the movie the temperature is kept at 37C, and images are taken once every 10 minutes. Exposure time is 10 ms for bright field and 500 ms for both mCherry and YFP.

Analysis of microscope movies is done using custom algorithms in ImageJ and MATLAB. For each frame, the phase image is converted to a binary mask of the cell colony. The mask and then used to find total mCherry and YFP fluorescence in the frame. After subtraction of background fluorescence, the total fluorescence is normalized by the total cell area (fluorescence intensity per pixel). For step induction experiments, fluorescence is normalized such that the maximum fluorescence of the lowest concentration induction is equal to 1 a.u. Figure S5 shows the microscopy analysis workflow. Error bars shown in microscopy time trace data are the standard error between analysis of different positions (n = 7 to 10) on the same experimental plate.

### Plate reader experiments

Plate reader data were collected on a Biotek H1MF machine using the kinetic read feature. Cells were incubated in the plate reader at 37C and shaken at 800rpm between reads. Cells were grown in clear bottomed 96-well microplates (PerkinElmer, ViewPlate, 6005182) and sealed with breathable clear membranes (Sigma Aldrich, Breath-Easy, Z380059). mCherry was read at excitation/emission of 580/610 with gain 140, Venus was read at 500/540 with gain 100.

Analysis of the data was done by taking fluorescence readings at late log phase for each independent well. Experimental conditions were done in triplicate and repeats were averaged. Fluorescence per OD was normalized by the fluorescence of a control strain (lacking mCherry or YFP) such that the cell autofluorescence equals 1 a.u.

### Model implementation

The model was implemented using the Simbiology toolbox in MATLAB and the ode15 solver (See Supplementary files for MATLAB code).

## Acknowledgements

VH is supported by the Department of Defense (DoD) through the National Defense Science & Engineering Graduate Fellowship (NDSEG) Program. Research supported in part by the Benjamin M. Rosen Bioengineering Center, the NIH/NRSA training grant 5 T3 2 GM07616, the Gordon and Betty Moore Foundation through Grant GBMF2809 to the Caltech Programmable Molecular Technology Initiative, and the Institute for Collaborative Biotechnologies through grant W911NF-09-0001 from the U.S. Army Research Office. The content of the information does not necessarily reflect the position or the policy of the Government, and no official endorsement should be inferred.

## Author Contributions

VH, ELCS, and WRW designed the research. VH and ELCS created the mathematical model and performed analysis. VH performed cell culture experiments and analysis. VH wrote the manuscript. WRW provided initial plasmids and critical technical insight.

